# *doepipeline*: a systematic approach to optimizing multi-level and multi-step data processing workflows

**DOI:** 10.1101/504050

**Authors:** Daniel Svensson, Rickard Sjögren, David Sundell, Andreas Sjödin, Johan Trygg

## Abstract

**Background:** Selecting the proper parameter settings for bioinformatic software tools is challenging. Not only will each parameter have an individual effect on the outcome, but there are also potential interaction effects between parameters. Both of these effects may be difficult to predict. To make the situation even more complex, multiple tools may be run in a sequential pipeline where the final output depends on the parameter configuration for each tool in the pipeline. Because of the complexity and difficulty of predicting outcomes, in practice parameters are often left at default settings or set based on personal or peer experience obtained in a trial and error fashion. To allow for the reliable and efficient selection of parameters for bioinformatic pipelines, a systematic approach is needed.

**Results:** We present *doepipeline*, a novel approach to optimizing bioinformatic software parameters, based on core concepts of the Design of Experiments methodology and recent advances in subset designs. Optimal parameter settings are first approximated in a screening phase using a subset design that efficiently spans the entire search space, then optimized in the subsequent phase using response surface designs and OLS modeling. *doepipeline* was used to optimize parameters in four use cases; 1) de-novo assembly, 2) scaffolding of a fragmented genome assembly, 3) k-mer taxonomic classification of Oxford Nanopore Technologies MinION reads, and 4) genetic variant calling. In all four cases, *doepipeline* found parameter settings that produced a better outcome with respect to the characteristic measured when compared to using default values. Our approach is implemented and available in the Python package *doepipeline*.

**Conclusions:** Our proposed methodology provides a systematic and robust framework for optimizing software parameter settings, in contrast to labor- and time-intensive manual parameter tweaking. Implementation in *doepipeline* makes our methodology accessible and user-friendly, and allows for automatic optimization of tools in a wide range of cases. The source code of *doepipeline* is available at https://github.com/clicumu/doepipeline and it can be installed through conda-forge.

## Background

Bioinformatic software tools frequently offer a number of outcome-related parameters for the user to set or change from their default values. These parameters may be different forms of input filters, or alter the behavior of the running algorithm. Parameters may be either quantitative or qualitative (multi-level) in nature. While it is advantageous to customize tools to a specific situation, it is not always obvious what effect changing parameters will have on the outcome. This may be due to lack of documentation, poor understanding of the algorithm, or interaction effects between parameters that are difficult to foresee. Additionally, software tools are commonly combined into pipelines, for example when calling genetic variants from raw sequence reads [1,2]. Pipelining tools in this manner further increases the complexity of selecting optimal parameter settings by increasing the numbers of both parameters and potential interaction effects. The settings for a particular data processing pipeline may also have to be tailored to the type of technology that was used to generate the data, for example the different platforms available for DNA sequencing which yield different error profiles [3]. In general, the strategy for selecting parameter settings therefore typically consists of using values derived from personal or peer experience and obtained in a trial-and-error fashion, or simply retaining the default values. This kind of non-systematic selection of parameter settings runs the risk of producing sub-optimal results.

The combined ranges of all possible parameter settings form a parameter space. To find the optimal point in the parameter space, an exhaustive brute-force search, commonly called a grid search, simply trying all possible combinations, is guaranteed to find the optimum. Since the number of combinations increases exponentially, exhaustive searching quickly becomes unfeasible as the number of parameters, and their ranges, grow. Instead, statistical Design of Experiments (DoE) can be used to span and investigate the parameter space in an efficient manner [4]. DoE aims to maximize information gain while minimizing the number of experiments required [5]. This is done by introducing variation into the system under investigation in a structured manner in order to explain how the parameters (*factors*) influence the result (*response*). This variation is introduced according to statistical designs for simultaneously varying the factor settings at a specific set of values (*levels*), and the system is modeled using statistical methods, for example with Ordinary Least Squares (OLS) regression [5–7]. The simplest type of statistical design is the full factorial design (FFD) where all combinations of factor levels are investigated in an exhaustive manner, meaning that they quickly become impracticable. To greatly reduce the number of experiments required, fractional factorial designs (FrFD) are used to investigate a structured subset of the FFD [6]. The problem is that FrFD are not trivial to use in situations where there are more than two levels to investigate, and that there is no obvious way to combine qualitative and quantitative variables. Recently, fractional factorial designs have been generalized into the so called generalized subset design (GSD) [8]. GSDs are balanced and near-orthogonal multi-level and multi-factor subset designs capable of mixing quantitative and qualitative factors, allowing for the investigation of a large and diverse set of parameters in an efficient manner. Compared to grid search, GSDs reduce the number of runs required to explore an equivalent parameter space by an integer factor, also called the *reduction factor*.

Although DoE is primarily used in analytical chemistry, a DoE approach has previously been applied by Eliasson et al to optimize software parameter settings in a liquid chromatography-mass spectrometry (LC-MS) metabolomics data processing pipeline [9]. In essence, this approach consists of sequentially updating a statistical design based on the predicted optimal configuration of settings, until they converge at an optimum. We build upon the approach proposed by Eliasson et al, and have developed a strategy for automated optimization of software parameter settings. We extend Eliasson et al’s approach with a screening phase using the recently developed GSD to efficiently span a much larger parameter space. We also make it possible to optimize multiple responses simultaneously. This extended approach may be used both for optimization of individual tools and for multiple tools organized into a pipeline. One crucial component is a well-defined objective function that you wish to minimize or maximize, i.e. there must be some way to objectively determine how well the pipeline is performing. Our strategy is software-agnostic and is implemented as a user-friendly Python package - *doepipeline*.

In this article, we outline our DoE-based strategy for a systematic approach to optimizing multi-level and multi-step data processing workflows, and exemplify the application of *doepipeline* with four cases; 1) de-novo assembly of a bacterial genome, 2) scaffolding of contiguous sequences (contigs) of a bacterial genome using 3rd generation sequencing (nanopore) data, 3) k-mer taxonomic classification of long noisy sequence reads generated by ONT MinION sequencing units, and 4) genetic variant calling in a human sample.

## Methods

We propose an approach for the optimization of software parameters, based on methods derived from statistical design of experiments. Our approach, which has been implemented in a python package (*doepipeline*), can be divided into two distinct phases:

1. Screening using a generalized subset design to find an approximate optimum. This phase also serves to find the best choice of categorical variables.
2. Iterative optimization, starting from the best point found by screening, based on the algorithm by Eliasson et al[9]. This phase optimizes only quantitative variables, meaning that categorical variables are fixed at the best values found during phase 1.

The screening and optimization phases are schematically illustrated in Figure 1 and described in more detail in the following subsections. Prior to screening and optimization, the user specifies what parameters to use as factors in the designs, whether they are categorical or numerical, and the permitted categories or value spans to be investigated. The user also specifies what process outcomes to use as response, and whether it should be maximized, minimized or reach a target value. In cases with several responses the user also needs to specify low/high limits and the target for each response. The responses are then re-scaled according to these limits and targets and combined into a single response using the geometric mean according to Derringer & Suich desirability functions [10]. In brief, when there are multiple responses each individual response is rescaled to be in the interval between 0 and 1, and it is 0 when outside accepted limits and 1 when better than the target. The rescaled responses are then combined into the overall desirability using the geometric mean.

**Figure 1.**
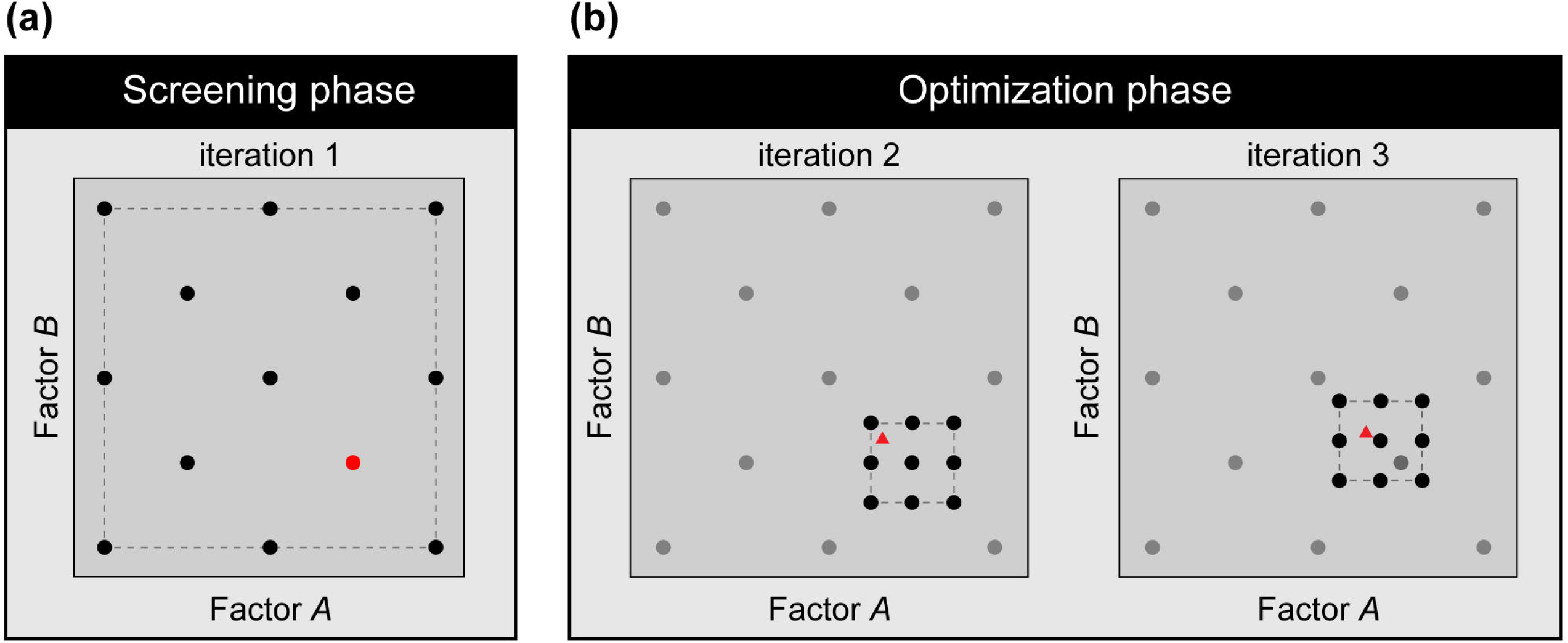
Schematic visualization of *doepipeline* design space movement. Example of optimization of two factors (A and B) through both the screening and the optimization phase, completed in 3 iterations. Each dot represents an executed pipeline with the parameters set by factors A and B. Triangles represent executed pipelines using the optima of an Ordinary Least Squares (OLS) model calculated in each optimization iteration. Red dots and triangles represent the best configuration of factors found in each iteration. Dashed lines represent the current high and low parameter settings in each iteration. *Screening phase*: a GSD using three levels and a reduction factor of 2 is used to span the design space. The pipelines are executed with the factor configurations suggested by the GSD and an approximate optimum is found (red dot). *Optimization phase*: in iteration 2, an optimization design is created around the best configuration found in the screening phase (black dots). In iteration 3, the design space is moved in the direction of the configuration of factors that produced the best result (red triangle) in iteration 2. *doepipeline* halts when the best response is produced by a configuration of factors that lies close to the center point (red triangle in iteration 3).

### Screening for Approximate Optimum

The purpose of the screening phase is to span the full search space to find regions with close to optimal performance. Screening is performed by executing the specified pipeline using combinations of factor configurations given by a GSD. Using a GSD effectively reduces the number of experiments to run, while optimally spanning the search space (Fig. 1a). The number of experiments required to investigate a given set of factors at a number of levels is approximately an integer fraction of the total number of possible combinations, which depends on the number of factors and their levels. A greater number of levels increases the resolution of the space searched during screening but also exponentially increases the number of runs required. We have found that five levels per numeric factor span large search spaces with a high enough resolution to give satisfactory results, but it is possible to set the number of levels individually for each factor in *doepipeline*. Similarly, we have found that the integer fraction of the full design that the GSD should represent can be safely set at the number of factors included in the design. However, this may also be controlled by the user by means of the *reduction factor* setting in *doepipeline*.

The screening phase also serves the purpose of setting the category to use for each categorical variable. For subsequent optimization, qualitative factors are fixed at the category of the best factor configuration according to the screening. By fixing qualitative factors, only numeric factors are investigated during the following optimization phase.

### Optimization of Numeric Factors

After selecting the best factor configuration during screening, numerical factors are optimized using response surface designs. The levels used in the screening design are here applied as anchor points for the new optimization design. A response surface design, for instance a central composite design, is constructed around the best configuration found. That is, the configuration of factor levels found to produce the best result during the screening phase is initially set as the center point in the new response surface design (Fig. 1b). If this configuration lies at the edge of a factor’s global design space (as defined by its min and max allowed values), the factor’s center point is shifted to the nearest screening level instead. This is done in order to keep the design within the global design space. After having set the center point for the new design, the high and low settings for each numeric factor are set to lie at the midpoints between the nearest screening levels respectively above and below the chosen center point, as indicated by the dashed line in Figure 1b. The span of each factor is then defined as the difference between the high and low settings. As during screening, the specified pipeline is executed using factor configurations given by the response surface design.

During each optimization iteration, pipeline performance is approximated using OLS regression [7]. By fitting a regression model the optimal configuration can be found by optimizing the response predicted according to the model. The factors included in the OLS model are selected either using a best subset approach or by using greedy forward selection; the latter is preferred when more than four factors are included in the design. If the predictive power (Q2) of the model is acceptable (Q2 > 0.5), the model is used to predict an optimal parameter configuration. Each numeric factor’s settings are then updated based on the best result in a manner similar to the algorithm given by Eliasson et al [9]. For each factor, the difference between the predicted best factor setting and the factor center point is calculated. If this distance is greater than 25% of the span of the factor, the high and low settings of the factor are updated in the direction of the best result. The default step length is 25% of the span of the factor, i.e. the high and low settings are moved 25% of the step length (Fig. 1b, iteration 3). We found that the algorithm did not always converge at this stage, but moved the design space back and forth between iterations. To alleviate this problem, we implemented design space shrinkage, which shrinks the design space span through multiplication by a so called *shrinkage factor* (typical value is 0.9, corresponding to 10 % shrinkage) between iterations, and found that it successfully improved convergence. If the proposed updated factor settings lie outside the predefined design space limits, the design is instead placed at the factor limits while keeping the same factor span. If the design has not moved between two iterations, or the best response is not improved upon compared to the previous iteration, the algorithm has converged and halts. If the optimization algorithm halts and responses have not reached their minimally acceptable values, the screening results are re-evaluated and a new optimization phase is run based on the results of the next best screening. At the end of the optimization iterations the factor configuration that has produced the best result throughout the iterations is chosen as the optimal configuration.

### Sequence data used in cases

The *Francisella tularensis sp. holarctica* strain FSC200 [11], and a genetic near neighbor *Francisella hispaniensis* strain FSC454 were chosen as an example dataset in case 1 to 3 of this study. The genome assembly of FSC200 is available as RefSeq assembly accesssion GCF_000168775.2 and genome assembly of FSC454 as RefSeq assembly accession GCF_001885235.1. Previously, generated Illumina HiSeq reads of FSC200 are available as NCBI SRA run SRR518502. This latter dataset was subsampled down to an estimated coverage of 100X (1.9M 100bp reads) for use in case 1, subsampling was performed with *seqtk* [12](v. 1.2-r94, installed through bioconda [13]).

New sequencing libraries were prepared from DNA extractions of the two bacterial strains using the SQK-LSK108 Ligation Sequencing Kit according to the manufacturer’s specifications and then sequenced using a FLO-MIN107 MinION flow cell (Oxford Nanopore Technologies, UK). MinION sequence reads for FSC200 are available as NCBI SRA run SRR9290761, and for FSC454 as NCBI SRA run SRR9290851. Subsampling down to 50 000 from 132 259 MinION reads for FSC200 and 15 000 from 15757 MinION reads for FSC454 was performed with a custom script, and the sequences were trimmed to a maximum length of 3000 bp as well as being sorted by length to increase classification speed.

### Case 1: de-novo assembly of a bacterial genome

In this example, we optimize the paired-end sequence assembler ABySS [14,15] (v. 2.0.2, installed through bioconda [13]) to assemble the genome of an isolate of *Francisella tularensis* ssp. *holarctica* (FSC200). ABySS has a total of 27 different parameters that can be specified by the user. Some are directly related to the running time and memory usage of the software (such as number of threads to use or bloom filter size), while others are related to the quality and/or characteristics of the resulting assembly (such as the size of k-mer or the minimum mean k-mer coverage of a unitig). For this example, we focused on the latter type of parameter. Hence, all parameters chosen to be part of the optimization were deemed to have a potential effect on the resulting assembly. The chosen parameters were: size of k-mer (*k*) (KMER), minimum mean k-mer coverage of a unitig (*c*) (MIKC), minimum alignment length of a read (*l*) (MIAL), and minimum number of pairs required for building contigs (*n*) (MIPA).

For this optimization we set the investigated factor space so that the default value for each factor was included within the span of each factors’ min and max values (Table 1). Although central to the ABySS algorithm, there is no default value for the k-mer size parameter. But since the value of the k-mer size is bounded by the actual read length it was still possible to define the GSD search space in a satisfactory way. For purposes of comparison, however, we considered a k-mer size of 31 to be the default setting. In this example we ran the initial screening with a reduction factor of 8, and used Central Composite Face-centered (CCF) designs in the following optimization iterations. We used a shrinkage factor of 0.9 (*-s*), and set the model selection method (*-m*) to greedy to speed up model selection. All other *doepipeline* settings were kept at default values.

**Table 1.**
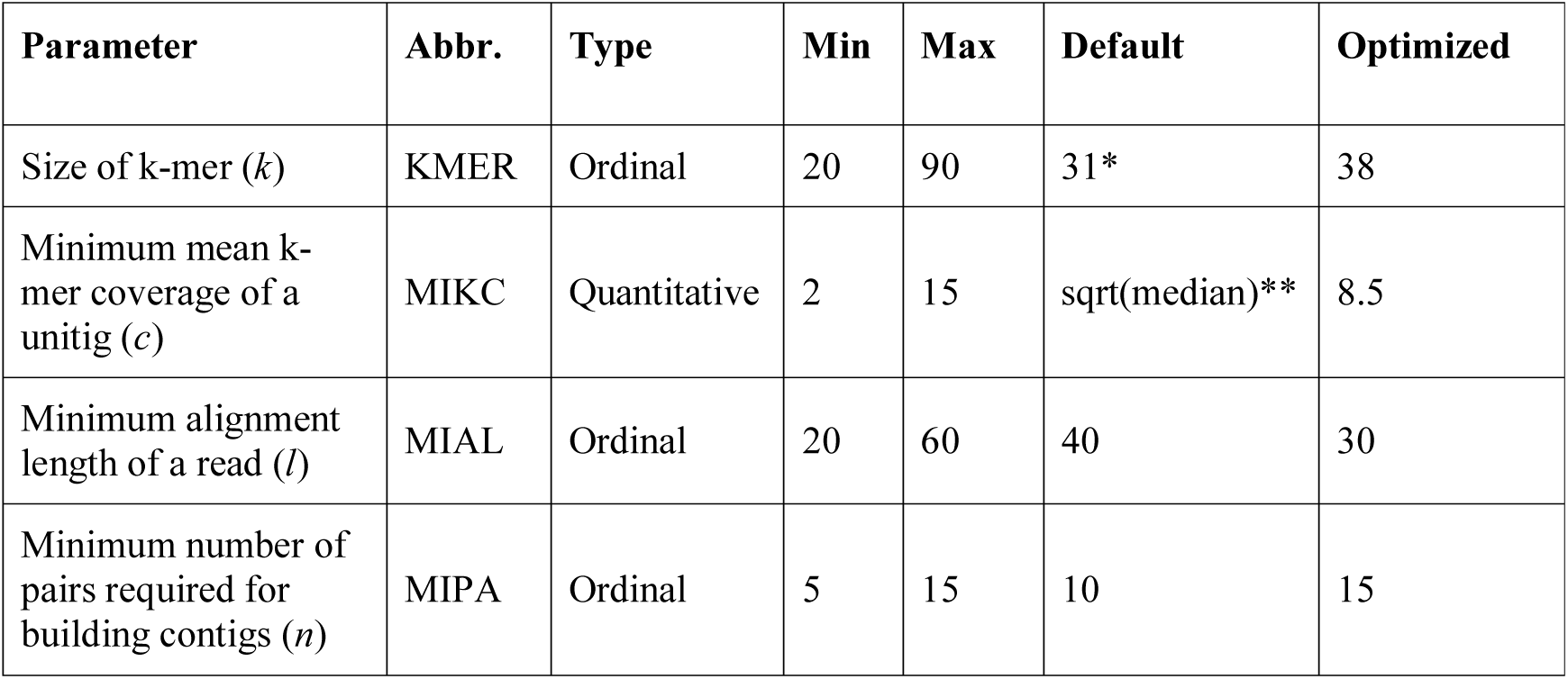
Factors in the de-novo assembly case. The four factors investigated in the de-novo assembly case are described below. The letter in parenthesis following the parameter name is the parameter used in the abyss-pe command line interface. Min and max values define the design space. *: There is no default value explicitly specified by the ABySS documentation. However here we used a k-mer size of 31 for comparison purposes. **: This refers to the square root of the median k-mer coverage, which is affected by the sequencing depth and choice of k-mer size. The optimized values are the combination of factor values that produced the best outcome, as found by *doepipeline*.

| Parameter | Abbr. | Type | Min | Max | Default | Optimized |
| --- | --- | --- | --- | --- | --- | --- |
| Size of k-mer (*k*) | KMER | Ordinal | 20 | 90 | 31* | 38 |
| Minimum mean k-mer coverage of a unitig (*c*) | MIKC | Quantitative | 2 | 15 | sqrt(median)** | 8.5 |
| Minimum alignment length of a read (*l*) | MIAL | Ordinal | 20 | 60 | 40 | 30 |
| Minimum number of pairs required for building contigs (*n*) | MIPA | Ordinal | 5 | 15 | 10 | 15 |

There are many metrics that can be used to evaluate the quality of a de-novo assembly, and which specific ones to use depends on what the assembly is to be used for [16,17]. Example metrics include the number of resulting contiguous sequences (contigs), the amount of total sequence covered by the assembly, and the N50 value. The latter is the length of the contig that, when the contigs are ordered by size, spans the midpoint of the total assembly. Hence, the N50 value can be viewed as an assessment of the quality of the assembly in terms of contiguity.

We used the total size of the assembly (tSeq), the number of contigs (nSeq), and the N50 value as responses. Since this optimization contained multiple responses, it was necessary to set low/high acceptable limits for each response, as well as target values to reach. The low and high limits for the responses were set with respect to the result obtained using the default settings with the same input data, meaning that the worst acceptable results are the default results. The target for the tSeq response was set to the reference genome size for FSC200 [11], while the targets for the nSeq and N50 responses were set to values that were considered achievable (Table 2).

**Table 2.**
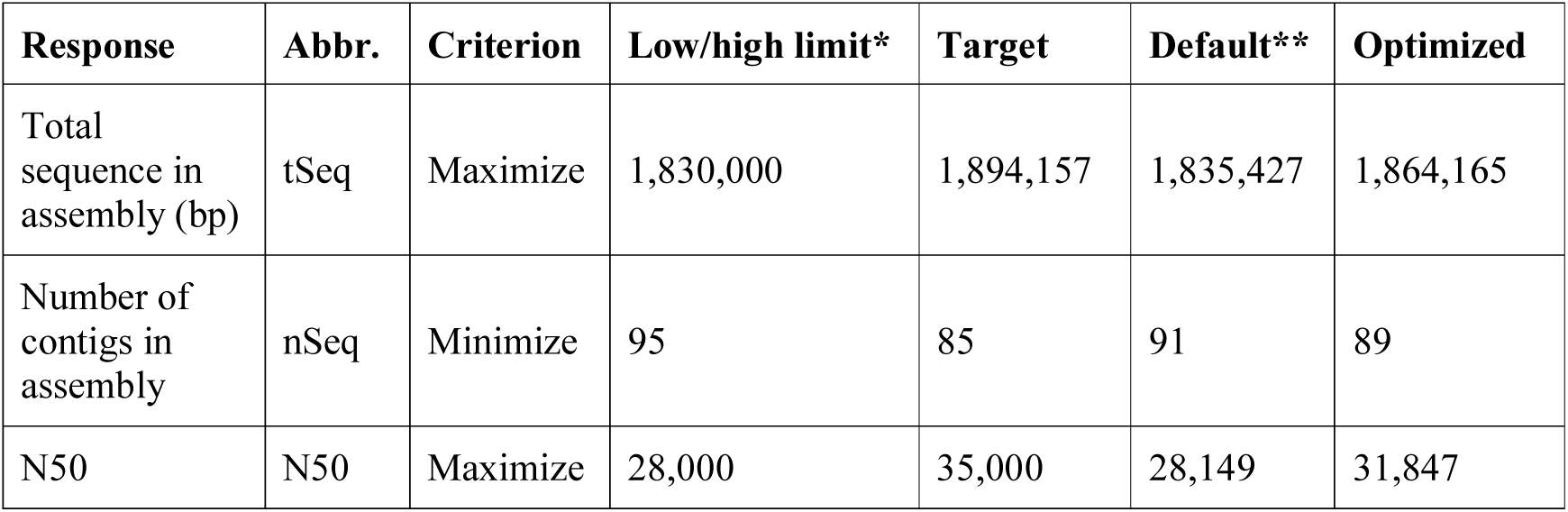
Responses in the de-novo assembly case. The three responses that were measured in the de-novo assembly case are described below. *: Responses that have the criterion maximize have a low limit, and those with the criterion minimize have a high limit. **: Default values are based on using a k-mer size of 31 and leaving all other parameters unchanged.

The data input to ABySS consisted of the subsampled Illumina HiSeq 2500 sequence data for FSC200 (see Sequence data used in cases). Prior to calculating the values for the responses we applied a length-based filter to the assembly using *Fastaq* [18] (v. 3.17.0), keeping only those contigs more than 1000 bp in length. This filter was also applied when calculating the response from the pipeline using the default parameter configuration. This is done because the very short contigs are typically made up of short repetitive sequences, and removing them simplifies the assembly graph and calculations on it. The software *seqstats* [19] was used to calculate the response values from the filtered assembly.

### Case 2: scaffolding of a bacterial genome assembly using long reads

Assembling a genome with short reads typically results in a fragmented assembly, consisting of a number of contigs. The way these contigs are connected with each other - in terms of ordering, distance, and direction - remains unknown. The reason for the fragmentation is that certain stretches of genomes have low complexity and are therefore impossible to resolve with short reads. One way of stitching together the contigs of an assembly is by using paired reads with long insert sizes, or - as is increasingly common - using long reads from, for example, the Nanopore or PacBio platforms. The long reads have an increased chance of spanning the low-complexity regions, effectively anchoring both ends of a pair of contigs together and thus resolving the gap. The process of connecting contigs together is referred to as scaffolding, and the resulting sequences are known as scaffolds.

SSPACE-LongRead [20] (SSPACE) uses long reads, such as those produced by the PacBio or Nanopore platforms, to scaffold an assembly. When running the software, the user can manipulate a total of six parameters that relate to the resulting scaffolds. We investigated whether manipulating some of the parameters would yield a better result than that achieved by running SSPACE (v. 1-1) with default parameter settings. We chose to optimize the minimum alignment length to allow a contig to be included for scaffolding (*a*) (ALEN), the minimum gap between two contigs (*g*) (GLEN), the maximum link ratio between the two best contig pairs (*r*) (RRAT), and the minimum identity of the alignment of the long reads to the contig sequences (*i*) (IDEN). As response, we maximized the N50 value of the resulting scaffolded assembly.

We set the investigated space for the factors so that the default value for each factor was included within the span of each factor’s min and max values (Table 3). For the optimization phase following the screening phase we chose to use a CCF design for the experiments. The reduction factor for the GSD was kept at the default value, i.e. the number of factors in the investigation, which in this case was 4. The model selection method (*-m*) was set to greedy and the shrinkage factor (*-s*) to 0.9. All other *doepipeline* settings were kept at default values.

**Table 3.** Factors in the scaffolding case. The four factors investigated in the scaffolding case are described below. The letter in parenthesis following the parameter name is the parameter used in the SSPACE command line interface. Min and max values define the design space. The optimized values are those that in combination produced the best outcome, as found by *doepipeline*.

| Parameter | Abbr. | Type | Min | Max | Default | Optimized |
| --- | --- | --- | --- | --- | --- | --- |
| Minimum alignment length to allow a contig to be included for scaffolding (*a*) | ALEN | Ordinal | 0 | 5000 | 0 | 0 |
| Minimum gap between two contigs (*g*) | GLEN | Ordinal | -3000 | 3000 | -200 | -750 |
| Maximum link ratio between two best contig pairs (*r*) | RRAT | Quantitative | 0.1 | 0.7 | 0.3 | 0.325 |
| Minimum identity of the alignment of the long reads to the contig sequences (*i*) | IDEN | Ordinal | 30 | 90 | 70 | 82 |

The input assembly had been constructed with ABySS [15] (v. 2.0.2) (k=71) and subjected to a contig length filter (>1000 bp). It consisted of 94 contigs between 1,685 and 87,479 bp in length, had an N50 of 27,549 bp, and totaled 1,800,912 bp prior to scaffolding. The assembly was constructed from the FSC200 Illumina HiSeq 2500 sequence data (see Sequence data used in cases). We include the assembly at the *doepipeline* github repository. The read set used for scaffolding consisted of 132,258 nanopore reads of between 163 and 108,214 bp in length (N50=679 bp), totaling 104,374,862 bp. *Seqstats* [19] was used to calculate the response from the scaffolded assembly.

### Case 3: k-mer classification

K-mer classification is a method used to assign taxonomic labels to short DNA sequence reads [21]. The method requires a precomputed database of k-mers generated from previously known and assembled genomes, for example all complete genomes in the NCBI database. When classifying a sample, the k-mer set of each read is calculated and compared with the database of known k-mers. The read is then assigned to the most specific taxonomic class within the database using the highest scoring k-mer root-to-leaf classification path following the taxonomic hierarchy. This method is implemented in, for example, the software package Kraken [22].

Kraken also uses a least common ancestor method, which re-classifies reads that are assigned to multiple taxonomic sub-classes under a parent node. A read with non-unique leaf assignment will then be assigned to the least common ancestor where there is little or no assignment conflict instead. The k-mer classification method implemented in Kraken can be applied to longer error-prone reads even though it is optimized for short accurate reads. However, it will be less accurate due to the different (higher) error frequencies and will therefore generate an increased rate of false positives.

In this study we used the software KrakenUniq [23] (v. 0.5.2). KrakenUniq builds upon the Kraken engine but additionally records the number of unique k-mers as well as coverage for each taxon. Three factors were used in the optimization: precision (PRES), minimum k-mer hits (MH) and a filter (FILT). We chose to use a CCF design in the optimization phase of *doepipeline*, the model selection method (*-m*) was set to greedy, and the shrinkage factor (*-s*) to 0.9. All other *doepipeline* settings were kept at default values. The F1 score (Eq. 1), which is the harmonic mean of precision and recall, was used as response.

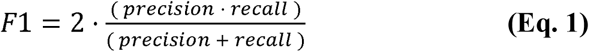

The input data were nanopore sequenced reads from two *Francisella* species, a target, *Francisella tularensis holarctica* (FSC200) and one near neighbor, *Francisella hispaniensis* (FSC454). The dataset was reduced to contain 50,000 *F. tularensis* reads and 15,000 (max) *F. holarctica* reads of maximum length 3000 bp, to increase the speed of classification and reduce potential bias (see Sequence data used in cases).

### Case 4: genetic variant calling

A single genetic difference with respect to a reference genome is referred to as a genetic variant, and the process of identifying these variants from sequence data is referred to as variant calling. Calling the simplest form of genetic variant, single nucleotide variants (SNV), from standard Illumina paired-end data is considered trivial nowadays, with F1 scores reaching 0.98 [24]. Because of this, we opted to optimize calling of short insertions and deletions (indels), which are slightly more complex and are harder to call correctly [24].

We used raw sequence data and high-confidence genetic (or “truth”) variant calls from a single well-studied individual, commonly known as NA12878. The raw sequence data (2×100 bp, 50X depth), which form part of the Illumina Platinum Genomes (PG) [25], were retrieved from the European Nucleotide Archive (ENA), study accession ERP001960 (run: ERR194147). The truth callset was a “hybrid” dataset, meaning it was produced by combining callsets obtained with different technologies and methodologies [25–27] as described in Krusche et al [28]. The truth set was downloaded from the PG GitHub repository [29].

The genome analysis toolkit (GATK) best practices workflow [1,2] was used as a guide for this variant calling case. Raw data processing was carried out in accordance with GATK best practices up to the point of having analysis ready reads, after which *doepipeline* was applied to optimize the variant calling and filtering steps. First, PICARD (v. 2.18.1) [30] was used to convert the sequence reads (FASTQ format) into unmapped BAM format (uBAM) and to mark Illumina adapters. We then mapped the reads to the hg19 reference (part of the GATK resource bundle) using BWA-MEM (v. 0.7.15 -r1140) [31,32] and marked duplicates using PICARD. Finally, Base Quality Score Recalibration (BQSR) was carried out using GATK (v. 3.8-1-0) [33] to obtain analysis-ready reads.

This case aimed to optimize variant calling and variant filtering, the remaining steps in the GATK best practices after obtaining analysis-ready reads. The calling was carried out using HaplotypeCaller, and the filtering was carried out using VariantFiltration, both tools within GATK. HaplotypeCaller offers around 20 adjustable parameters while the VariantFiltration tool expects custom-specified cutoffs for annotations in the variant call format (VCF) file. GATK suggests four annotations by which to filter indels. In order to include a meaningful number of parameters at each step we chose to optimize the two steps sequentially. Performing sequential optimization allowed us to investigate 4 parameters for each step, 8 in total. We first optimized the calling step while keeping the parameters in the filtering step at their default settings. We then optimized the parameters for the filtering step using the output from the highest scoring experiment in the first step. For the calling step we chose to optimize the global assumed mismapping rate for reads (*globalMAPQ*) (GMQ), the minimum base quality for calling (*mbq*) (MBQ), the minimum reads per alignment start (*minReadsPerAlignStart*) (RAS), and the minimum confidence threshold for calling (*stand_call_conf*) (SCC). For the filtering step we chose to optimize the quality by depth (*QD*) (QD), the read position rank sum test (*ReadPosRankSum*) (RPRS), the Fisher test for strand bias (*FS*) (FS), and the strand odds ratio (*SOR*) (SOR). To further reduce the size of the optimization, we chose to optimize only against variants on chromosome 1. However, we screened for any overfitting of the parameters by executing the variant calling and filtering pipeline across all autosomes and chromosome X with the optimized parameters.

The following settings were used for both optimizations. We set the space investigated for the factors so that the default value for each factor was included within the span of each factor’s min and max values (Table 5). The design of choice for the screening phase was the CCF design. The reduction factor for the GSD was increased to 8, reducing the number of experiments. The model selection method (*-m*) was set to greedy and the shrinkage factor (*-s*) to 0.9. All other *doepipeline* settings were kept at default values.

Performance metrics and tools to assess the accuracy of variant calling in a standardized manner are crucial, and the benchmarking team of the Global Alliance for Genomics and Health (GA4GH) have made significant progress with respect to this [28]. The GA4GH benchmarking team has developed a benchmarking tool, hap.py [34], that can compare a high-confidence (or “truth”) variant callset with a user-made single-sample callset, also known as the query callset, and output performance metrics. For a certain set of confident regions (specified by BED file), concordant variants in the two callsets should be considered true positives (TP), while discordant variants should be considered either false positives (FP) or false negatives (FN) depending on which callset they appear in. Hap.py also outputs the F1 score (see case 3 methods) for variants passing the VCF filters, which was used as the response in this case.

### Grid search comparison

We compared the results from *doepipeline* to those from grid search, which is a common methodology for optimizing parameters. Grid search is done by evaluating the parameter performance for all possible combinations of parameter settings, the so-called parameter grid. For the comparison to be relevant, we performed the grid search at the same resolution as the GSD screening step in each case. In other words, we tested all possible combinations of the factor setting levels (typically 5 levels per factor).

#### doepipeline

##### Implementation

*doepipeline* is fully implemented in the Python programming language and source code is available for download at github (https://github.com/clicumu/doepipeline) and installable with conda-forge [35,36] and through PyPi. Generation of statistical designs is carried out through the python package PyDOE2 [37], in which the GSD has been implemented.

##### Usage

Configuration of the optimization is done in a structured YAML file with sections for the experimental design and for the pipeline steps (commands) to run. The design section includes the names of the factors investigated and their min/max values (design space), the responses and their goals (minimize/maximize), and the type of design to use in the optimization phase. The pipeline section is where each individual pipeline step is specified. In each iteration, *doepipeline* takes the pipeline steps as configured and substitutes the parameters under investigation with the values given by the statistical design. A batch script is created for each pipeline step, with any parameter values substituted, and the execution of it is controlled by *doepipeline*. Pipeline steps are executed either in parallel mode, where all experiments are run at the same time, or in sequential mode where each pipeline with all of the steps is executed in sequence. For reference, we provide example YAML files at the github repository.

Today, scientific data processing can include vast amounts of data and/or require substantial computing power. In such cases, data processing is commonly performed on compute clusters that typically use some queueing system in order to handle all user requests for resources. An example of such a queueing system is the Slurm Workload Manager [38] (Slurm). To accommodate users of compute clusters, we have implemented Slurm support for *doepipeline*. If using Slurm, specify the Slurm options in the YAML file as you would when running a regular Slurm job. The Slurm options are transferred by *doepipeline* to the batch script which is then submitted to Slurm using *sbatch*.

After optimization, the parameter values suggested by *doepipeline* are saved in the working directory for the optimization. Additionally, there is a rich log file that can be investigated to follow the workflow.

## Results

### Case 1: de-novo assembly of a bacterial genome

The goal of de-novo assembly is to combine raw sequence reads into a representation of an organism’s genome, i.e. to obtain as contiguous a genomic sequence as possible. Due to the characteristics of the genome sequence itself, in combination with short reads, this process can be difficult. For example, sequence reads from less complex segments of the genome will map to more than one position, causing ambiguities that are not possible to resolve, and this in turn leads to fragmentation of the assembly.

One popular sequence assembler is ABySS [15], which provides 27 different user-controlled parameters. We set up an example for optimization of de-novo assembly software parameters using ABySS (see Methods section). *doepipeline* ran for two iterations before halting. Thus, the best response was obtained in the first iteration, in the GSD screening phase. The experimental sheet and corresponding response values from the GSD screening and iteration 2 are included as Additional file 1. Using the optimized parameter settings (Table 1), we obtained a 1.6% and 13.1% increase in the investigated responses tSeq and N50, and a 2.2% reduction in nSeq as compared to when abyss-pe was run with default settings (Table 2). Optimizing the parameters using the grid search option required 625 experiments to be run, and it resulted in the same combination of parameter settings as when using *doepipeline* (see Additional file 2 for grid search result). By comparison, doepipeline required 97 experiments to be run.

### Case 2: scaffolding of a bacterial genome assembly using long reads

Scaffolding is the process of connecting together contigs obtained from an assembly step. In this example we aimed to optimize parameters for the scaffolding software package SSPACE-LongRead [20], which relies on long reads to span the low-complexity regions that are typically found between the contigs of an assembly. *doepipeline* ran for three iterations before halting, obtaining the best result in the second iteration. The response values and parameter settings investigated in each iteration are included in Additional file 3. The response (N50) value obtained when using the default parameter settings was 1,141,889 bp. Using the optimized parameter settings (Table 3) resulted in a 66.9% increase in the response (1,905,883 bp). Optimizing the parameters using the grid search option required 625 experiments to be run, compared to 211 experiments using *doepipeline*, and it resulted in a best N50 value of 1,868,309, which is slightly lower than the result obtained using *doepipeline* (see Additional file 4 for grid search result).

### Case 3: k-mer classification

K-mer classification is used to gather information about the species content of a metagenomic sample. It is possible to visualize the general distribution of species through the reads classified or to identify the presence/absence of reads classified to specific targets. By using third generation sequencing techniques, such as Oxford Nanopore, it is possible to classify reads from an unknown sample in real time. But due to the long error-prone reads produced by third-generation sequencing machines, there is a greater risk of misclassification. At the genus level this is not usually a problem. But when it comes to discriminating between pathogenic and non-pathogenic species, misclassification may become problematic; in particular false positive signals of pathogenic species may be obtained. We investigated the KrakenUniq [23] (v. 0.5.2) algorithm and used *doepipeline* to find optimized settings for long error-prone reads in order to increase the ratio of true positives to false positives using the F1-score as response. KrakenUniq also has a filter that may reduce the number of false positive reads. The filter will adjust each assigned read up the tree until the desired threshold is met, where the threshold is the number of assigned k-mers divided by the number of unique k-mers in that category [23].

Optimization ran for three iterations before halting and the best results were found during the second iteration. The experimental sheet and corresponding response values from the GSD screening and optimization iterations are included in Additional file 5. Using the optimized parameter settings (Table 4), we were able to increase the F1 score by 0.065% from 0.993690 to 0.994341, compared to when running KrakenUniq with default settings.

**Table 4.** Factors in the k-mer classification case. The three factors investigated in the k-mer case are described below. Min and max values define the design space. The optimized values are those that in combination produced the best outcome, as found by *doepipeline*. *: The KrakenUniq documentation to our knowledge does not state what the default value is.

| Parameter | Abbr. | Type | Min | Max | Default | Optimized |
| --- | --- | --- | --- | --- | --- | --- |
| Minimum k-mer hits | MH | Ordinal | 1 | 200 | * | 14 |
| Standard deviation of the relative errors of the estimate | PRES | Ordinal | 10 | 18 | 12 | 17 |
| Minimum tax-ID score threshold | FILT | Quantitative | 0 | 0.05 | 0 | 0 |

Optimizing the parameters using the grid search option required 125 experiments to be run, compared to 76 experiments using *doepipeline.* The grid search resulted in a best F1 score of 0.994169 which is slightly lower than the result obtained using *doepipeline* (see Additional file 6 for grid search result).

### Case 4: genetic variant calling

Variant calling is the process of determining genetic variants (or mutations) from genetic sequence data. In this case we aimed to find optimized parameters for a widely used variant calling framework, the genome analysis toolkit (GATK). Specifically, we sequentially optimized two of the steps carried out by GATK: variant calling and variant filtering (see methods).

In the optimization for the first step (variant calling), *doepipeline* ran for three iterations before halting, obtaining the best result in the second iteration (F1=0.9707). In the optimization for the second step (variant filtration), *doepipeline* ran for four iterations before halting. The best result (F1=0.9716) was obtained in the fourth iteration when the optimum predicted by the model was validated. This optimum was too far from the design space edges for *doepipeline* to move the design space and initiate another iteration, and thus it halted execution. The response values and parameter settings investigated in each iteration are included in Additional file 7 (variant calling step) and 8 (variant filtering step). The included parameters and their default and optimized settings are listed in Table 5.

**Table 5.**
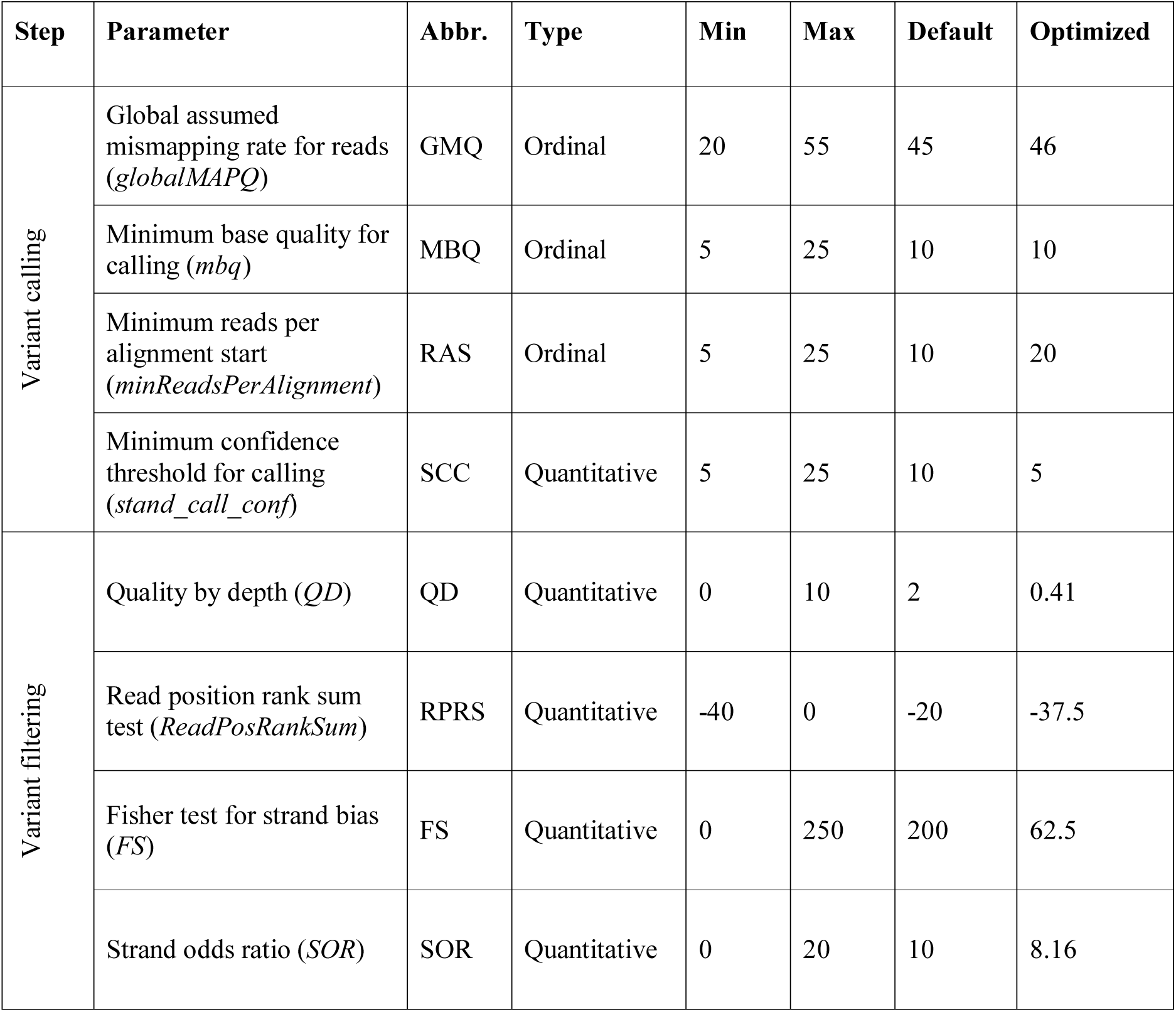
Factors in the variant calling case. The factors investigated in the variant calling case are described below. The optimization was carried out sequentially for two main steps, variant calling and variant filtering, and which step each factor belongs to is indicated. For the variant calling step, the factor’s corresponding command line flag is given in parentheses after the parameter name. For the variant filtering step, the corresponding information tag annotated in the VCF file is indicated in parentheses. The min and max values define the design space. The default values for all factors are also indicated; for the calling step they are the build-in default values of the HaplotypeCaller tool, while for the filtering step the default values are those recommended by the GATK team. The optimized values are those that in combination produced the best outcome, as found by *doepipeline*.

| Step | Parameter | Abbr. | Type | Min | Max | Default | Optimized |
| --- | --- | --- | --- | --- | --- | --- | --- |
| Variant calling | Global assumed mismapping rate for reads (*globalMAPQ*) | GMQ | Ordinal | 20 | 55 | 45 | 46 |
|  | Minimum base quality for calling (*mbq*) | MBQ | Ordinal | 5 | 25 | 10 | 10 |
|  | Minimum reads per alignment start (*minReadsPerAlignment*) | RAS | Ordinal | 5 | 25 | 10 | 20 |
|  | Minimum confidence threshold for calling (*stand_call_conf*) | SCC | Quantitative | 5 | 25 | 10 | 5 |
| Variant filtering | Quality by depth (*QD*) | QD | Quantitative | 0 | 10 | 2 | 0.41 |
|  | Read position rank sum test (*ReadPosRankSum*) | RPRS | Quantitative | -40 | 0 | -20 | -37.5 |
|  | Fisher test for strand bias (*FS*) | FS | Quantitative | 0 | 250 | 200 | 62.5 |
|  | Strand odds ratio (*SOR*) | SOR | Quantitative | 0 | 20 | 10 | 8.16 |

As the optimization was performed only on chromosome 1, we wanted to see how well the optimized parameter settings carried over into a variant calling and filtering pipeline applied across all autosomes and chromosome X. This analysis resulted in an F1 score of 0.9713, while using the default settings resulted in an F1 score of 0.9702.

Optimizing the parameters using the grid search option resulted in a best F1 score of 0.9715, which is marginally lower than the results obtained using *doepipeline*. Five experiments in the first step of the grid search optimization (calling) resulted in the same highest F1 score (see Additional file 9). We therefore ran five parallel instances of the second step of optimization (filtering) using the different VCF files from the five best experiments in the first step. This inflated the number of required experiments from the expected 1250 to 3750 experiments in total, compared to 280 experiments with *doepipeline*. The five parallel optimizations of step two all yielded the same set of 12 combinations of settings producing an equally high F1 score (0.9715) (see Additional file 10). Validation across all autosomes and chromosome X using all 60 combinations of parameter settings (5 times 12) yielded a best F1 score of 0.9712, again marginally lower than for doepipeline.

## Discussion

Selecting parameter settings for a data processing pipeline is complex, since the influence of the parameters on the end result is not always obvious. While the value of personal and peer experience should not be underestimated, our approach provides a systematic way of determining optimal settings. Specialized tools to optimize particular bioinformatic software tools have been proposed previously. For example, VelvetOptimizer [39] can be used to optimize the k-mer and coverage cutoff parameters of the Velvet assembler [40] and KmerGenie can be used to make an informed decision on the choice of k-mer in de Bruijn based assemblers [41]. However, a generalized, software-agnostic optimization approach is preferable, especially when several tools are used together in a pipeline.

Here we present such a generalized strategy for automated sequential optimization of software parameters, employing core concepts of DoE methodology. We have implemented our strategy in a user-friendly python package, *doepipeline*. The optimization strategy and the use of *doepipeline* was exemplified in four bioinformatics use cases; de-novo assembly of a bacterial genome using Illumina reads, scaffolding a bacterial genome assembly using nanopore reads, k-mer classification of metagenomic third generation sequencing data, and genetic variant calling. In all four cases, we saw an improvement in the measured response variables as compared to when using the default parameter settings. The improvement of the measured responses in our examples ranged between 0.065% and 66.9%. We compared the results from *doepipeline* to results from standard grid searches, and *doepipeline* achieved equally good or better results using significantly lower numbers of evaluations/experiments. Grid search is typically limited to running a single optimization phase evaluating all points in the parameter grid with no further refinement. This is in contrast to *doepipeline*, which is adaptive and refines the parameter settings based on the best results from the previous phase, allowing it to find better performing parameter settings than grid search.

One of the advantages of our proposed strategy is the use of a GSD in a screening phase prior to the optimization phase. Compared to Eliasson et al[9], we are able to screen a much larger design space efficiently prior to optimization using the GSD-based approach. In order for the optimization phase to converge in a feasible number of iterations, the design space should be restricted in some way. Deciding the range of each of the factors without guidance risks creating too narrow or wide a design space. Instead, the screening allows the user to set up a relaxed (wide) design space in which to investigate and to approximate the optimal factor combination. The approximation represents a substantiated initial center point around which to set up a narrower optimization design. The screening phase will also identify promising values for any qualitative factors and fix them before optimization. Thus, the GSD screening phase can be viewed as a systematic approach to restricting the design space for the subsequent optimization phase. Similar results can be achieved using stochastic optimization methods such as random search [42], commonly applied within the machine learning community. Random search can effectively reduce the number of runs required, but the final results are probabilistic and may not be optimal, depending on each particular random draw. By using structured space-filling designs, *doepipeline* deliberately spans more of the search space rather than relying on randomness. We note that the multi-phase workflow of *doepipeline* has conceptual similarities to Bayesian hyperparameter optimization [43], in refining the parameter choice based on promising parameter regions from earlier iterations. However, *doepipeline* uses statistical designs that are guaranteed to fill the parameter space and structured refinement around promising points rather than randomly sampling promising regions with higher probability.

The fraction of the full design that a GSD represents can be controlled with the reduction factor parameter in *doepipeline*. We ran the optimization of ABySS (case 1) with a GSD reduction factor of 8, but another optimization of ABySS where a reduction factor of 10 was used produced the same response values (data not shown) in fewer experimental runs (45 as opposed to 70). In addition, there was a degree of overlap among the response values in the GSD iterations (Additional files 1, 3, and 5). Overall, this could indicate that it is meaningful to try running the GSD with a higher reduction factor than the recommended default, and/or reducing the number of levels, further reducing the number of experiments.

Currently, *doepipeline* leverages cloud computing capability through the Slurm workload managing system. Given the recent development and consolidation of workflow managing systems [44] it would be possible to integrate *doepipeline* with for example SnakeMake [45] or NextFlow [46], similar to other implementations [47,48].

During development and testing of *doepipeline* we saw the design space moving back and forth between iterations in the optimization phase. We hypothesized that this behavior was because either the underlying function was not modeled properly or the function was flat within the investigated design space. To counteract this phenomenon we implemented three features; i) no prediction of the optimal factor combination if the predictive power (Q2) of the model was low (default: Q2<0.5), ii) validation of the predicted optimal factor combination by carrying out the pipeline with those factor settings, and iii) shrinking the span of the factors between iterations. After implementing these three features, *doepipeline* consistently converged to satisfactory results.

Specifying the pipeline in a YAML file allows for flexible configurations of commands to be run, essentially enabling optimization of any pipeline run on the command line. However, the number of parameters will typically increase with the length of the pipeline under investigation. At the same time there is a soft constraint on the number of parameters that can be investigated simultaneously. This constraint will be related to the problem currently under investigation and depends on the computational complexity of the pipeline, and on the available computational and time resources. Instead of doing a global optimization of parameters, i.e. optimizing the entire pipeline at once, an alternative approach is to run sequential optimizations in which only a section of the pipeline at a time is optimized while keeping the default parameter values for the rest of the pipeline [9]. This type of sequential optimization is not yet fully implemented in *doepipeline* and is a feature for future updates. Sequential optimization of a pipeline currently requires that an optimization is carried out for each step of the pipeline and that the optimized parameter values so obtained are manually updated for the subsequent steps of the pipeline.

## Conclusion

Our proposed strategy represents a systematic approach to the optimization of software parameters. Our implementation in the software-agnostic and user-friendly package *doepipeline* could potentially serve as a starting point for experimenters and bioinformaticians who currently rely on default settings or common practice when running their data processing pipelines.

## Declarations

### Ethics approval and consent to participate

Not applicable

### Availability of data and material

The datasets generated and analyzed during the current study (cases 1-3) are available in NCBI BioProject accessions PRJNA510899 (SRA run: SRR9290761), PRJNA16087 (SRA run: SRR518502), PRJNA548675 (SRA run: SRR9290851). The assembly used as input to case 2 is available at the *doepipeline* github (https://github.com/clicumu/doepipeline). The sequence data used for case 4 is available at the ENA with study accession ERP001960 (run: ERR194147), and the truth callset along with confident regions BED file is available at the Platinum Genomes github repository [29]. The GATK bundle is available at the GATK FTP ftp://ftp.broadinstitute.org/bundle/hg19/.

### Consent for publication

Not applicable

### Authors’ contributions

DSv, AS and JT initiated the study, DSv and RS designed the algorithm and implemented the python code. DSv and DSu analyzed and interpreted the four example cases. All authors wrote, read and approved the final manuscript.

## Acknowledgements

This study makes use of sequence data generated at the Swedish Defence Research Agency by Edvin Karlsson and Emelie Samuelsson-Näslund. The authors would like to acknowledge support from the Science for Life Laboratory (SciLifeLab) and the National Genomics Infrastructure (NGI) for providing assistance in massively parallel sequencing. Computations were carried out at the Uppsala Multidisciplinary Center for Advanced Computational Science (UPPMAX) and the High Performance Computing Center North (HPC2N), part of the Swedish National Infrastructure for Computing (SNIC). The authors thank Matilda Rentoft and Mattias Eliasson for fruitful discussions.

## Additional material

**Additional file 1**

Excel file (.xlsx).

Complete experimental sheets for case 1, *doepipeline* iterations 1 and 2. Contains factor settings and response values for all experiments in these iterations.

**Additional file 2**

Excel file (.xlsx).

Complete experimental sheet for case 1, grid search. Contains factor settings and response values for all executed experiments.

**Additional file 3**

Excel file (.xlsx).

Complete experimental sheets for case 2, *doepipeline* iterations 1, 2, and 3. Contains factor settings and response values for all experiments in these iterations.

**Additional file 4**

Excel file (.xlsx).

Complete experimental sheets for case 2, grid search. Contains factor settings and response values for all executed experiments.

**Additional file 5**

Excel file (.xlsx).

Complete experimental sheets for case 3, *doepipeline* iterations 1, 2, and 3. Contains factor settings and response values for all experiments in these iterations.

**Additional file 6**

Excel file (.xlsx).

Complete experimental sheet for case 3, grid search. Contains factor settings and response values for all executed experiments.

**Additional file 7**

Excel file (.xlsx).

Complete experimental sheets for case 4, step 1, *doepipeline* iterations 1, 2, and 3. Contains factor settings and response values for all experiments in these iterations.

**Additional file 8**

Excel file (.xlsx).

Complete experimental sheets for case 4, step 2, *doepipeline* iterations 1, 2, 3, and 4. Contains factor settings and response values for all experiments in these iterations.

**Additional file 9**

Excel file (.xlsx).

Complete experimental sheet for case 4, step 1, grid search. Contains factor settings and response values for all executed experiments.

**Additional file 10**

Excel file (.xlsx).

Complete experimental sheet for case 4, step 2, grid search. Contains factor settings and response values for all executed experiments.

